# Relaxin-3 neurons of the zebrafish periaqueductal gray mediate behavioral responses evoked by darkness

**DOI:** 10.64898/2026.09.26.754586

**Authors:** Joanne Shu Ming Chia, Michael Martin, Grace M Padilla, John Jefferson, Venkatesh Krishna, Gilles Vanwalleghem, Ruey-Kuang Cheng, Seetha Krishnan, Matthew Lovett-Barron, Tod R Thiele, Suresh Jesuthasan

**Author notes:** Corresponding Author: SJ. Co-first authors (Joanne Shu Ming Chia, Michael Martin). Co-senior authors (Matthew Lovett-Barron, Tod R Thiele, Suresh Jesuthasan).

## Abstract

Ambient illumination influences many aspects of brain function, including responses to threats. One brain region proposed to mediate these effects is the periaqueductal gray (PAG), an evolutionarily conserved midbrain structure that contains multiple subpopulations of peptidergic neurons. Here we ask which cell types in the larval zebrafish PAG respond to changes in ambient illumination. By registering *post hoc* gene expression to neurons recorded during exposure to sensory stimuli, we show that GABAergic PAG neurons expressing the neuropeptide Relaxin-3 (*rln3a*+) are persistently activated by darkness, while a neighboring subpopulation of GABAergic PAG neurons expressing *penka* is not. Zebrafish lacking *rln3a* show a reduction in locomotion during darkness. We propose that darkness recruits Relaxin-3-releasing PAG neurons to promote vigilance and potentiate defense circuits. These results identify a specific peptidergic population of PAG neurons that are potentially relevant to behavioral state dysregulation associated with excessive darkness.

## Introduction

The midbrain periaqueductal gray (PAG) is a critical hub in defensive behavior, pain modulation, and arousal state, through its regulation of motor and neuromodulatory systems. The PAG in mammals has classically been organized into columns^1^, with glutamatergic neurons of the dorsal PAG controlling active defensive responses including escape^2–4^, while GABAergic neurons in the ventrolateral region (vlPAG) act locally to regulate passive responses including freezing^3^ and analgesia^5^. Recently, however, a transcriptomic study has shown the existence of numerous subpopulations within the PAG, which can be classified based on neuropeptide expression^6^. A subpopulation of cells expressing the peptide cholecystokinin (CCK) responds to distant threats and controls escape, distinct from the freezing induced by broadly stimulating the surrounding neurons^7^. Thus, understanding how the PAG regulates brain state and behavior requires a consideration of subpopulations with distinct neurochemical properties, beyond the canonical columnar/spatial organization.

Among the neuropeptides produced by PAG neurons are Enkephalin^8^ and Relaxin3^9^. While the role of Enkephalin in pain modulation is well established^10–13^, the function of Relaxin3 is less clear. Global knockout experiments and receptor manipulation in mice suggest that Relaxin3 influences diverse processes including locomotion, arousal, stress responses, and feeding^14–16^. However, because the *relaxin3* gene is strongly expressed in the nucleus incertus as well as the PAG^17^, it is unclear what the function of the PAG subpopulation is. Here, we use zebrafish larvae, which allow for cellular-resolution functional imaging in the intact animal, to investigate the sensory response properties of *rln3a*-expressing cells in the PAG.

Primarily, we investigate the functional responses of cells expressing the *relaxin3a* gene (*rln3a*) to changes in ambient illumination. In diurnal animals, light reduces fear behavior^18^ and pain perception^19^, while darkness has the opposite effect. In mice, which are nocturnal, light increases fear responses^20^. The diurnal zebrafish provides a useful experimental model to understand the effects of ambient illumination on brain state in other diurnal vertebrates, including humans, where excessive darkness is associated with mood disorders^21^.

## Results

### *rln3a* and *penka* are expressed in adjacent GABAergic neurons in periaqueductal territory in larval zebrafish

Relaxin3-expressing neurons have previously been identified in the larval zebrafish midbrain and proposed to correspond to the PAG based on anatomical position, as well as regional overlap with expression of *penka*^22,23^. Using whole-mount double fluorescent *in situ* hybridization (FISH) in 6-7 dpf larvae followed by volumetric confocal imaging, we found that both *rln3a+* and *penka+* neurons flank the cerebral aqueduct in the midbrain tegmentum, immediately ventral to the deep layers of the optic tectum (Figure 1A, B; Supplemental Movie 1, 2). The two populations were largely intermingled, with *penka*+ cell bodies extending further along the ventrocaudal axis; spatial overlap between transcripts was rare, with only one or two double-positive neurons per fish. Both populations were predominantly GABAergic, with 82.8 ± 5.4% of *rln3a+* neurons (consistent with prior transgenic characterization^23^) and 90.9 ± 2.7% of *penka+* neurons expressing *gad1b/gad2*, and little glutamatergic co-expression in either population (Figure 1D-F). These results suggest that *rln3a* and *penka* are expressed in different subsets of GABAergic neurons in the larval zebrafish PAG.

**Figure 1.**
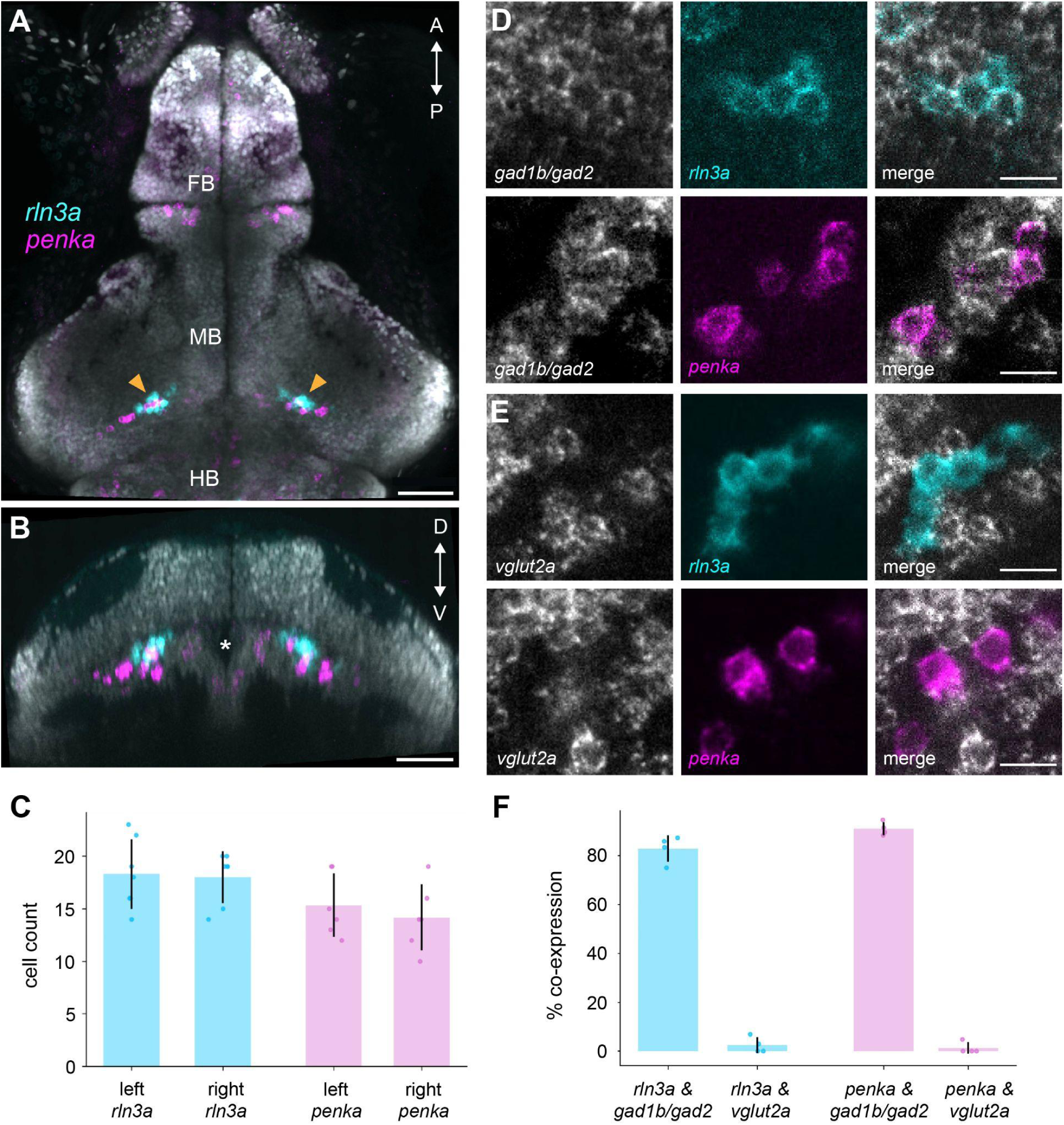
*rln3a* and *penka* neurons are closely situated inhibitory cell populations in the zebrafish PAG. (**A, B**) Dorsal (**A**) and transverse (**B**) maximum-intensity projections of a 6 dpf larval zebrafish brain showing wholemount double FISH for *rln3a* (cyan) and *penka* (magenta) counterstained with DAPI (gray). Arrowheads indicate the bilateral PAG clusters; the asterisk indicates the ventricle. FB, forebrain; MB, midbrain; HB, hindbrain; A, anterior; P, posterior; D, dorsal; V, ventral. Scale bars, 50 μm. (**C**) Cell counts for left and right *rln3a* and *penka* populations (mean ± SD; individual animalsshown as dots; n = 5 fish). (**D**) Representative confocal images showing co-expression of *rln3a* (cyan) and *penka* (magenta) with the GABAergic markers *gad1b/gad2* and (**E**) the glutamatergic marker *vglut2a* (gray). Scale bars, 5 μm. (**F**) Percentage of *rln3a* and *penka* neurons co-expressing GABAergic (*gad1b/gad2*) or glutamatergic (*vglut2a*) markers (mean ± SD; n = 4 fish).

### *rln3a*+ neurons are activated by darkness

Given that bright light inhibits GABAergic neurons in the ventrolateral PAG in mice^24^, the region expressing *rln3*^9^, we hypothesized that *rln3a*+ neurons in zebrafish are responsive to visual input. We first examined the responses of pan-neuronal populations in the larval zebrafish PAG, using two-photon calcium imaging of *Tg(elavl3:GCaMP6f)* larvae exposed to discrete pulses of widefield light. Blue light was used in initial experiments, as this wavelength strongly activates ipRGCs that are implicated in the regulation of brain state by light^25–27^. Neurons in the midbrain tegmentum showed distinct responses to light and darkness, with OFF responses located laterally and ON responses located medially (Figure 2A, B, C; n = 8 fish). The lateral cells consistently showed a reduction in GCaMP6f fluorescence in the presence of light and an increase in darkness (Figure 2C and Supplemental Figure 1A; Pearson coefficient ± standard deviation = 0.62 ± 0.16; n = 123 cells); responses in the medial cells were less correlated (Pearson coefficient = -0.32 ±0.27; n = 85 cells). We next examined the effects of different wavelengths of light, to assess whether PAG responses were driven by cone photoreceptors. While red, blue, and violet light consistently elicited a reduction in GCaMP6f fluorescence in lateral neurons, the response to darkness varied with wavelength (Supplemental Figure 1B, C). No consistent illumination-correlated response was detected in medial neurons (Supplemental Figure 1D). In contrast to the responses in the lateral PAG, we did not detect illumination-or darkness-induced activity in the nucleus incertus (Figure 2C). Together, these results indicate that the lateral PAG of zebrafish larvae contains a subset of neurons whose activity increases in darkness, potentially involving a retinal OFF pathway.

**Figure 2.**
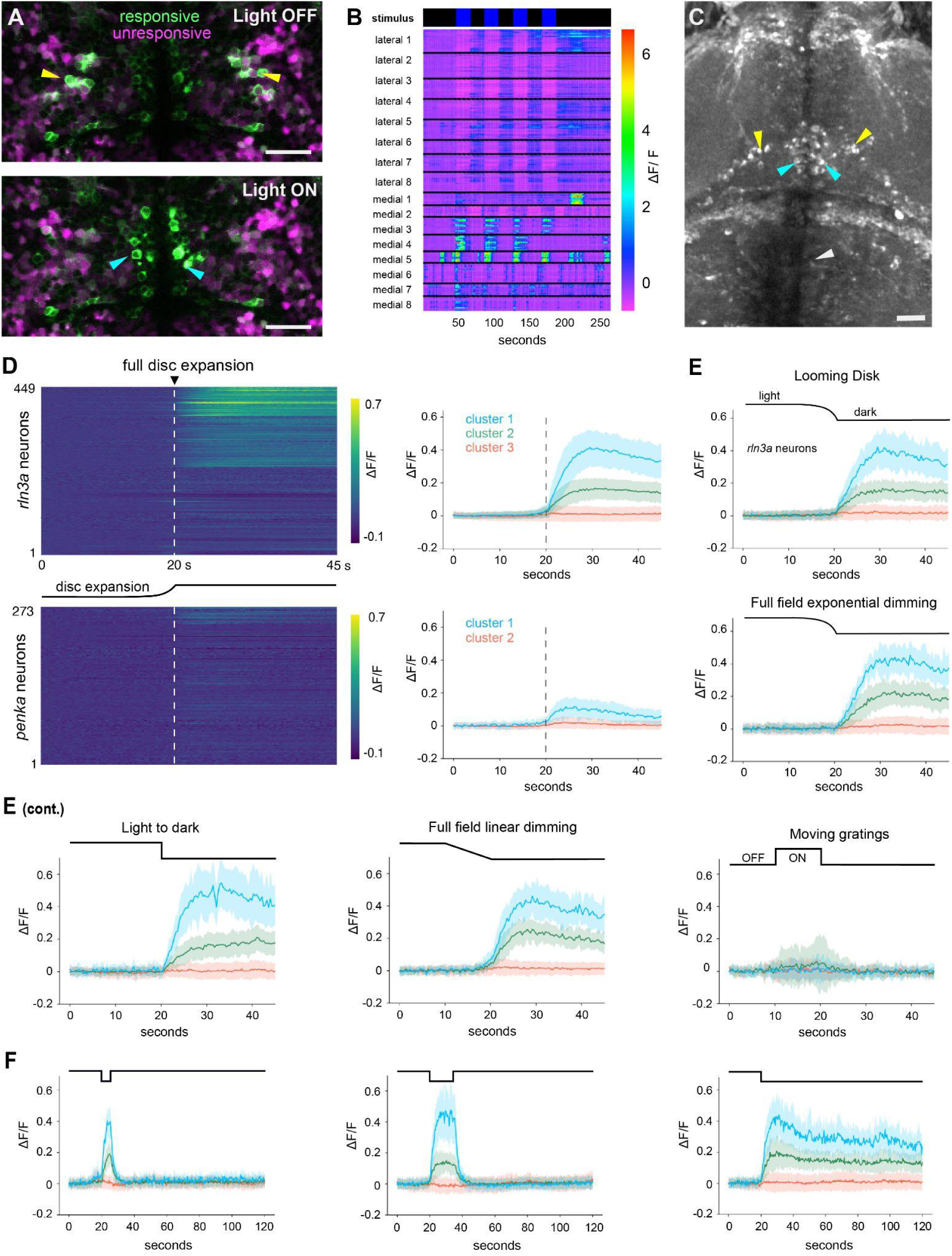
*rln3a* neurons in the PAG are selectively activated by darkness. (**A**) Midbrain response of one 7 dpf *Tg(elavl3:GCaMP6f, gad1b:DsRed)* larva to four pulses of blue light. Cells in the lateral region of the PAG (A top image; yellow arrowheads) have elevated GCaMP6f fluorescence in darkness, while cells in the medial tegmentum (**A** bottom image: cyan arrowheads) occasionally show increase in the presence of light. (**B**) Heatmap showing changes in GCaMP6f fluorescence in medial and lateral PAG cells, in 8 larvae. The fish in panel A is number 4 in this rastermap. (**C**) A standard deviation projection of a multi-plane time series of an 8 dpf *Tg(elavl3:GCaMP6f)* larva exposed to 3 pulses of violet light and 3 pulses of blue light. Activity can be seen in the lateral (yellow arrowhead) and medial (cyan arrowheads) PAG, but not in the nucleus incertus (white arrowhead). (**D**) Responses of all *rln3a* neurons (top; n = 449 across 13 fish) and *penka* neurons (bottom; n = 273 across 11 fish) during looming disc presentation, shown as heatmaps of ΔF/F sorted by k-means cluster assignment (left) and mean ΔF/F ± SD per cluster (right). Dashed line indicates full disc expansion at 20 s. (**E**) Mean ΔF/F ± SD for *rln3a* neuron clusters in the same 4 fish, each presented with a looming disc, full-field exponential dimming, an abrupt light-to-dark transition, full-field linear dimming, and moving gratings (OFF/ON indicated). Stimulus profile shown above each plot. (**F**) Mean ΔF/F ± SD for *rln3a* clusters across dark periods of varying duration, illustrating sustained activity during darkness and abrupt termination upon light return (n = 3 fish). Scale bars = 25 µm (**A, C**); larvae are shown in dorsal view, with anterior to the top.

The darkness-responsive neurons in the lateral PAG were GABAergic, as judged by fluorescent protein expression under the *gad1b* promoter (Figure 2A; n = 8 fish). To determine whether darkness-driven PAG neurons expressed *rln3a* and/or *penka*, we combined calcium imaging with *post-hoc* gene expression labeling, registering imaging volumes to *in situ* hybridization data at single-cell resolution^28,29^ (Supplemental Figure 2). We used a looming disc (Supplemental Figure 3), which simulates an approaching predator through progressive expansion and ends in sustained darkness, thereby allowing an assessment of whether the PAG response occurs to a gradual dimming or to darkness. Neurons classified as expressing *rln3a* displayed prolonged increases in activity upon full expansion of the disc (Figure 2D left panel). K-means clustering of identified neurons recorded across 13 animals identified three response clusters (Figure 2D right panel): a high-amplitude group (CL1: 17.1%, 77 of 449 neurons), an intermediate group (CL2: 30.1%, 135 of 449), and an unresponsive group (CL3: 52.8%, 237 of 449), with a mean of 16 *rln3a*+ neurons per fish responding. The magnitude of calcium responses for *rln3a+* cells ipsilateral and contralateral to the stimulus were similar (Supplemental Figure 4). By contrast, *penka*+ neurons showed minimal recruitment to these visual stimuli, with only 8.4% of neurons displaying a low-amplitude response, and the remainder largely unresponsive (Figure 2D).

Given that we only saw *rln3a*+ neural activity after full disc expansion, we directly tested whether gradual reduction in illumination would activate *rln3a*+ neurons, by exposing larvae to whole-field exponential dimming or whole-field linear dimming (n = 4). We observed increases in neuronal activity upon the onset of darkness (Figure 2E), but not during a gradual linear dimming (Figure 2E); forward-moving gratings elicited little to no response (Figure 2E). The darkness-responsive subset of *rln3a*+ neurons maintained their elevated fluorescence throughout prolonged darkness, with an initial steep rise followed by sustained activity; this activity ceased as soon as luminance increased (Figure 2F). These results demonstrate that the activity of *rln3a*-expressing neurons is correlated with complete darkness, rather than partial dimming or looming patterns. Furthermore, the sharp drop in activity at the offset of darkness indicates that these neurons’ persistent activity is driven by constant sensory input (i.e.,of darkness, or of the absence of widefield light).

While we found that *rln3a*+ neurons were responsive to darkness, the intermingled population of *penka*+ neurons was minimally responsive. However, we reasoned that these neurons may be active in other contexts. For instance, enkephalinergic neurons of the mammalian PAG have a long-established role in descending antinociception, forming the cellular substrate through which opioids gate pain suppression^30^. We therefore asked whether *penka*+ neurons in the zebrafish PAG are responsive to nociceptive stimuli, rather than visual stimuli. We presented zebrafish with head-targeted thermal stimuli^31^ (Supplemental Figure 3) at three intensities, while imaging both *penka*+ and *rln3a*+ neurons, identified by post-imaging *in situ* hybridization and volume registration. Low (7 mW of 980nm laser) and medium (28 mW) heat levels produced little tail movement and no clear changes in the activity of *penka*+ neurons. However, we found that the highest intensity of heat (65 mW) evoked robust escape behavior, and strongly recruited *penka+* neurons; approximately 26% of these cells were grouped into the most active response cluster (Figure 3A-D). This selective recruitment at noxious heat levels is consistent with a conserved antinociceptive role for these neuron types, and adds further support to the homology between this region of the zebrafish midbrain and the mammalian vlPAG. We note that *rln3a*-expressing neurons were also recruited at the highest heat intensity, suggesting they respond broadly to aversive and arousal-promoting stimuli. Thus, subclasses of GABAergic neurons in the zebrafish PAG, distinguished by their unique expression of neuropeptide genes, differ in their functional responses to sensory stimuli. Notably, *rln3a*+ neurons are distinguished by their persistent engagement by darkness, suggesting the Relaxin-3 neuropeptide may play a unique role in the behavioral response to darkness.

**Figure 3.**
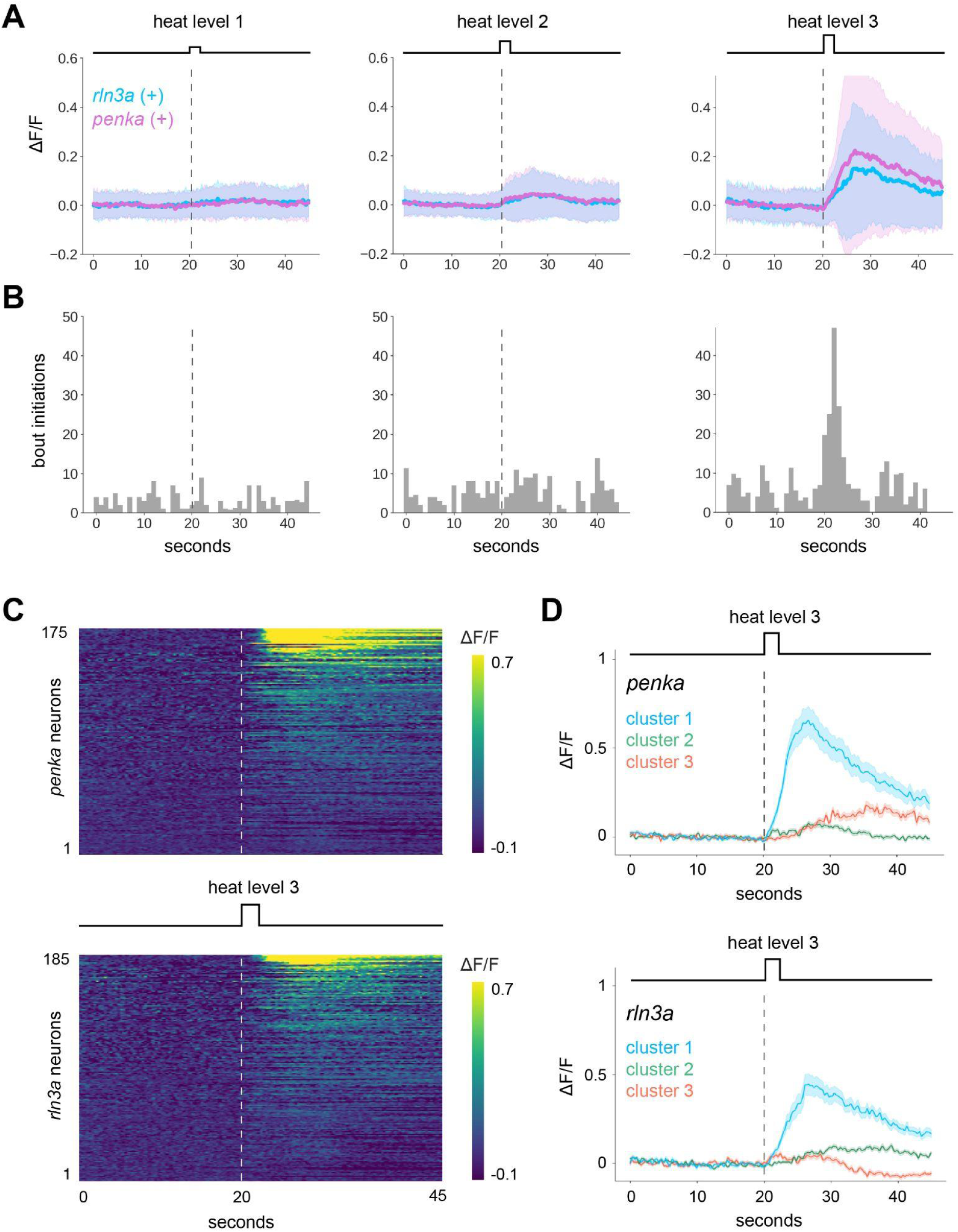
*penka* and *rln3a* neurons are recruited by noxious heat at escape-evoking intensities. **(A)** Mean ΔF/F ± SD for all *penka* (magenta) and *rln3a* (cyan) neurons during application of graded noxious heat at three intensities (heat levels 1–3; n = 6 fish). Dashed line indicates stimulus onset at 20 s. **(B)** Histograms of bout initiations per 1-s bin across the same stimulus epochs, illustrating the increase in escape behavior at heat level 3. **(C)** Heatmaps of ΔF/F for *penka* (top, n = 175) and *rln3a* (bottom, n = 185) neurons during heat level 3, sorted by cluster assignment. **(D)** Mean ΔF/F ± SD per cluster for *penka* (top) and *rln3a* (bottom) neurons during heat level 3. Dashed line indicates stimulus onset.

The persistent recruitment of *rln3a*+ neurons by darkness raised the question of whether their activity relates to defensive behaviors that darkness evokes. Following full expansion of the looming disc, larvae either initiated swimming bouts or remained immobile during the post-expansion window. Separating trials by the presence of bouts (Supplemental Figure 5A), we found that both responsive clusters (CL1, CL2) were recruited on trials with and without bouts (Supplemental Figure 5B), but that peak responses (Max ΔF/F) were modestly larger on trials in which larvae remained immobile than on trials in which they produced tail movements, on hemispheres both ipsilateral and contralateral to the looming stimulus (Supplemental Figure 5C). The activity of *rln3a*+ neurons is therefore elevated in association with the freezing response to darkness on a seconds-long timescale. Importantly, this acute correlation need not predict the direction of the phenotype produced by chronic loss of the peptide as Relaxin3 signals through the G-protein-coupled receptor RXFP3 (Relaxin-family peptide receptor 3), and neuromodulation of this kind alters postsynaptic excitability over seconds-to-minutes rather than at the speed of fast synaptic transmission^32^. Sustained activity of *rln3a*+ neurons during darkness may thus be positioned to exert a slow influence on sensory integration and locomotor drive across long time courses, rather than directly gating individual swim bouts.

### Loss of *rln3a* alters behavioral response to darkness

To test the behavioral function of *rln3a*, we generated a mutant using CRISPR/Cas9. The Relaxin3a protein consists of two chains, A and B (Figure 4A, B), which are generated by cleavage of the propeptide. We designed a guide RNA to a region upstream of the receptor binding residues in chain B (Figure 4B), with the aim of generating a mutation that prevents binding of the ligand to its receptor. Sequence analysis of the offspring of embryos injected with this guide RNA and Cas9 showed a 7-base pair deletion at the target site in one line (Figure 4C). This results in a frameshift that is predicted to give rise to a truncated protein lacking the highly conserved receptor-binding motif. Given the presence of a premature stop codon, transcripts of the *rln3a* gene in the *rln3a^sq4sj^* mutant are predicted to undergo nonsense-mediated decay^33^. Using real time quantitative RT-PCR of whole adult fish brains, we found expression levels of *rln3a* in the homozygous mutant (ΔCt M = -9.31, 95% CI [-9.69, -8.93]) were 16-fold lower than the WT siblings (ΔCt M = -5.32, 95% CI [-5.65, -4.99]), showing a depletion of the mRNA after the mutagenesis (Figure 4D; Supplemental Figure 6).

**Figure 4.**
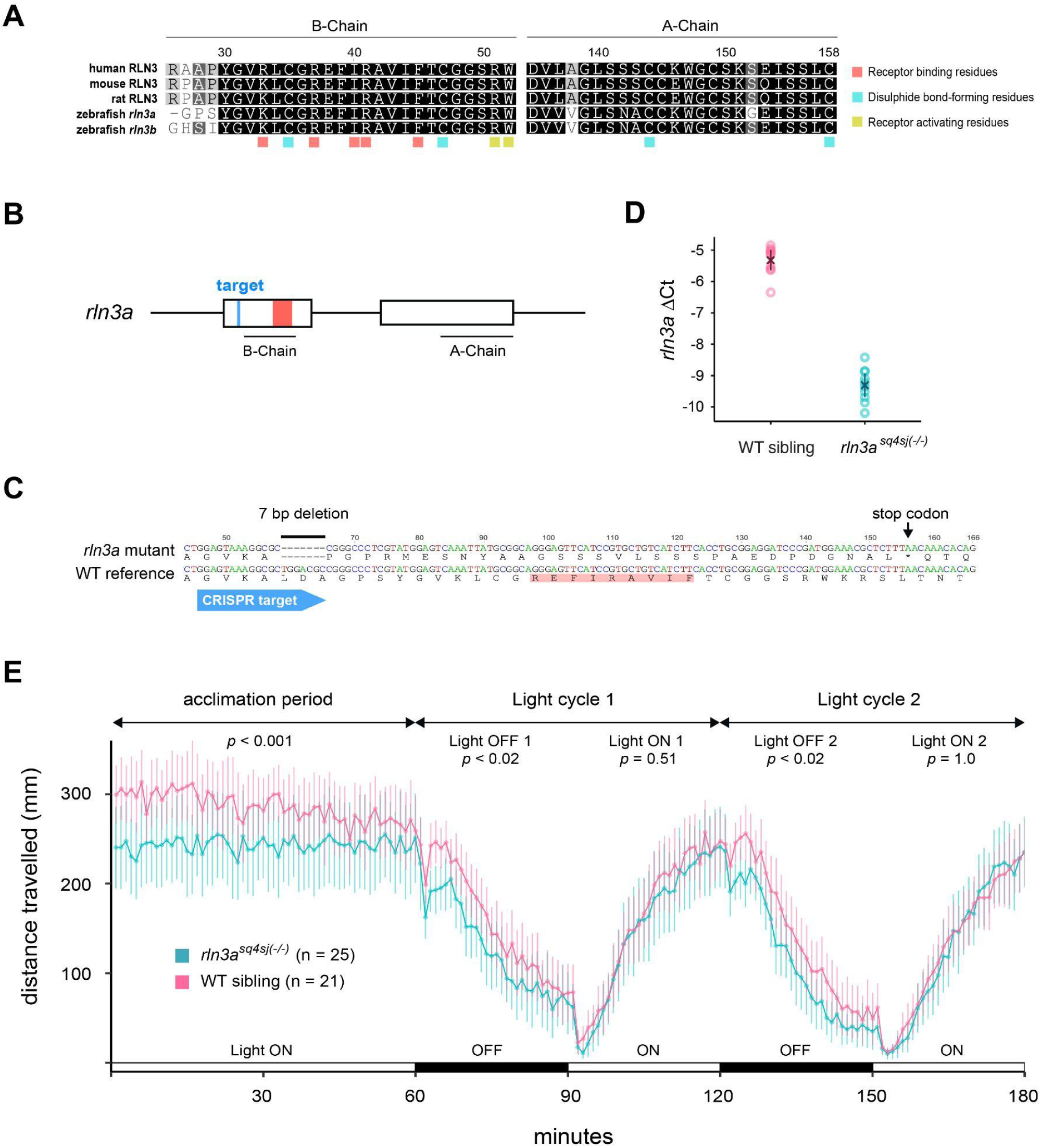
*rln3a* influences darkness-evoked locomotion. (**A)** Alignment of zebrafish Relaxin 3a and Relaxin 3b protein sequences with the Relaxin 3 in human, mouse and rat. The residues that contribute to structure and function, which are highlighted in colored boxes, are highly conserved. The B-chain contains receptor-binding residues, which are highlighted in red. Cysteine residues, which contribute to the formation of disulphide bonds and exist in both domains, are highlighted in cyan. Residues that contribute to receptor activation upon ligand binding are highlighted in yellow. (**B**) Schematic of the *rln3a* locus. The CRISPR target is within the first exon and upstream of the receptor binding region. (**C**) Nucleotide alignment between wild type and mutant sequence with the predicted translated peptide sequence. The *rln3a^sq4sj^* mutant has a 7 bp deletion; as a result, the receptor-binding sites are no longer present and a premature stop codon has formed. (**D**) Expression levels of *rln3a* in mutant and WT sibling fish (n = 10 samples per group),shown as ΔCt (Ct_ref_ – Ct_gene_). Unfilled circles represent individual fish; crosses indicate means with 95% CI. (**E**) Mean distance travelled per minute by *rln3a^sq4sj^* mutants (n = 25) and wild-type siblings (n = 21) during a 60-minute acclimation period in light, followed by two cycles of 30 minutes of darkness and 30 minutes of light. Error bars show 95% confidence intervals. *p* values are Bonferroni-corrected pairwise contrasts between genotypes within each epoch, from a linear mixed model (see Methods).

To assess the effect of this mutation on darkness-driven behaviors, we measured the change in larval zebrafish swimming behavior that is initiated by sudden darkness^34^. Following a 60-minute baseline habituation period, larval zebrafish mutants and wild-type siblings were exposed to cycles of alternating 30-minute light and dark periods. During the baseline period, *rln3a* mutants were hypoactive relative to siblings (*p* < 0.001;*post-hoc* Bonferroni-corrected contrasts), possibly reflecting a reduced arousal response to placement in the assay dish, with activity in the two groups converging toward the end of the habituation period (Figure 4E). Both groups decreased activity during dark periods and increased activity in light, but *rln3a* mutants moved significantly less than siblings during dark cycles and during the baseline habituation period, while no difference was detected during light cycles (linear mixed model, genotype x epoch interaction: F(4, 8138) = 21.0, *p* < 0.001; *post-hoc* Bonferroni-corrected contrasts: baseline *p* < 0.001, dark cycle 1 *p* = 0.018, dark cycle 2 *p* = 0.016, light cycles *p* > 0.5; Figure 4E). The absence of a detectable genotype difference during the light phases of both light/dark cycles indicates that the reduced activity in darkness is not simply a continuation of the early baseline hypoactivity. Although Relaxin3 signaling is eliminated in both the PAG and nucleus incertus in these mutants, the absence of darkness-evoked activity in the nucleus incertus neurons (see Figure 2C) suggests the dark-phase behavioral phenotype is a consequence of the loss of Relaxin3 signaling from darkness-activated PAG neurons.

## Discussion

In this study, we examined the activity of two peptidergic cell types in the zebrafish midbrain, which we posit are homologous to cell types in GABAergic regions of the mammalian PAG. Using anatomical analyses, behavioral studies, combined functional imaging with *post hoc* molecular identification, and analysis of a neuropeptide mutant, we have discovered that *rln3a*+ neurons, but not neighboring *penka*+ neurons, are strongly activated by darkness. In contrast, strong nociceptive thermal stimuli recruit broad sets of midbrain cells that encompass both peptidergic populations. These results suggest that peptidergic populations for threat detection are conserved in the zebrafish midbrain, with cell type-specific roles in the response to luminance and thermal nociception.

Several lines of evidence support the designation of the *rln3a* and *penka* populations as residing within the zebrafish homolog of the vlPAG. Firstly, both populations flank the cerebral aqueduct in the ventral midbrain tegmentum, a defining cytoarchitectural feature of the PAG across vertebrates^35^. Secondly, each is predominantly GABAergic, consistent with the substantial inhibitory neuron content of the mammalian PAG^6^. Thirdly, their spatial intermingling parallels the co-distribution of Relaxin3- and Enkephalin-expressing neurons that are enriched in ventral lateral regions of the mammalian PAG^7,9^. Lastly, selective recruitment of *penka* neurons following noxious heat, but not visual stimuli, parallels the antinociceptive profile of enkephalin-expressing vlPAG neurons in mammals^30,36^, providing functional support for this homology. While the term “griseum centrale” has been suggested to describe the homolog of the PAG in zebrafish^37,38^, we have opted to use the terminology of the PAG here, to reflect the anatomical and functional similarity with the mammalian midbrain structure that regulates a variety of instinctive and threat-directed behaviors^35^

We found that neurons expressing *rln3a* were unresponsive to moving gratings or during gradual dimming, and instead responded at the end of dimming (i.e. to darkness) and maintained elevated activity throughout the period of darkness. This pattern raises the possibility that *rln3a*-expressing neurons in the zebrafish are driven by the retinal OFF pathway, potentially by a subset of low-sensitivity populations^39^. Our observation that the response is affected by wavelength suggests the involvement of retinal cones, and is consistent with the finding that movement induced by darkness is influenced by wavelength of the preceding light^40^. The absence of darkness responses in nucleus incertus *rln3a*+ neurons confirms that this luminance sensitivity is specific to the PAG population of neurons producing this neuropeptide, and supports the interpretation that the dark-phase behavioral phenotype of *rln3a* mutants is caused by loss of Relaxin3 signaling in the PAG specifically. The magnitude of the behavioral reduction in *rln3a* mutants likely underestimates the full contribution of Relaxin3 signaling, since the *rln3a* paralogue *rln3b* is also expressed in the PAG and could partially compensate for the loss of *rln3a*^41^. Zebrafish PAG neurons expressing *rln3a* have broad projection targets including the medulla, diencephalon, hypothalamus, and optic tectum^23^a profile consistent with a broadcast-like neuromodulatory projection neuron^42^ rather than a locally-projecting interneuron^42^. Projections of *rln3a*+ cells to the hypothalamus are of particular interest, given that hypothalamic CRF neurons are critical to the aversive effects of darkness^43^.

A substantial body of work has identified the effects of illumination levels on brain activity, including circuits affecting threat response, focusing on the role of ipRGCs^44^, which respond to the presence of light. In humans, for example, fMRI indicates that light inhibits the amygdala^45^. The data here suggest that the retinal OFF pathway may also signal to the PAG. An interesting next step would be to determine whether darkness elicits a similar response in the human brain.

## Methods

### Fluorescent in situ hybridization

HCR probes, hairpins, and buffers were purchased from Molecular Instruments. HCR processing followed a previously published protocol^29^: zebrafish were fixed overnight in 4% paraformaldehyde in 1× PBST at 4 °C. After washing (3 times in 1× PBST, 5 min each), larvae were permeabilized for 10 min in 100% (v/v) methanol at −20 °C and then rehydrated (50% (v/v) methanol, then 25% (v/v) methanol, then 2× SSCT; 5 min each). Hybridization with split probes was performed overnight in 2× SSCT, 10% (v/v) dextran sulfate, 10% (v/v) formamide at a probe concentration of 4 nM. The next day, larvae were washed (3 times in 2× SSCT, 30% (v/v) formamide at 37 °C, then 2 times in 2× SSCT at room temperature; 20 min each) and then incubated in amplification buffer (5× SSCT, 10% (v/v) dextran sulfate). During this time, dye-conjugated hairpins were heated to 95 °C for 1 min and then snap-cooled on ice. Hairpin amplification was performed by incubating individual larvae in 50 μl of amplification buffer with B1, B3, and B5 hairpins at concentrations of 240 nM overnight in the dark. Samples were washed 3 times in 5× SSCT for 20 min each, then mounted in 2–3% low-melting-point agarose, covered in 2× SSCT or PBS, and immediately imaged on a Zeiss LSM 800 AxioExaminer confocal microscope using a W Plan-APO 20x 1.0 NA objective.

### Functional and anatomical imaging

For experiments in Singapore (Figure 2A-D, Supplemental Figure 1), zebrafish larvae (aged 5 - 14 dpf) were incubated in mivacurium and then embedded in low-melting temperature agarose (1.2-2.0 % in E3) in a glass-bottom dish (Mat Tek). They were imaged on a Nikon two-photon microscope (A1RMP; Tokyo, Japan), attached to a fixed stage upright microscope, using a 25x water immersion objective (NA = 1.1). The femtosecond laser (Coherent Vision II; Santa Clara, CA) was tuned to 920 nm. Fish were imaged during the day only, between 10 am and 7 pm. The facility lights were on from 8 am to 10 pm. Fish were selected at random for imaging. Imaging was carried out according to protocols approved by the Biological Resources Centre IACUC (number 120771).

Light stimuli were generated by 5 mm blue (458 nm peak emission), violet (405 nm) and red (650 nm) LEDs. They were powered by a 5 V TTL signal from a control computer and synchronized with image capture using a National Instruments DAQ board, controlled by the Nikon Elements software. Light intensity at the sample was 0.13 mW/cm^2^.

To analyse calcium imaging data, cells were segmented using CellPose (https://www.cellpose.org), using cpsam_v2, with a diameter setting of 6, flow threshold = 0 and cell probability threshold = -1.0. Suite2p^46^ was used to identify cells with activity. The Multi-Cell Analysis plugin for FIJI^47^ was then used to generate DeltaF/F data, which was plotted using Graphpad Prism or Matlab..

*University of Toronto experiments (Figure 1, 2, Supplemental Figure 2):* Zebrafish were raised and bred at 28 °C on a 14 h light/10 h dark cycle. Two-photon calcium imaging experiments were performed using 6-8 dpf *Tg(elavl3:H2B:GCaMP6f)^jf7^* larvae. The evening before experiments, fish were embedded in 2% low-melting agarose. Neuronal activity was collected on a Scientifica resonant scanning two-photon microscope using an Olympus 25x NA 1.05 water immersion objective (XLPLN25XWMP2). A Spectra-Physics MaiTai Ti:Sapphire laser tuned to 920 nm was used for excitation. GCaMP6f fluorescence was collected from a volume containing the PAG region using 8-10 planes spaced 7 µm apart at 512x512 pixel resolution, at a rate of ∼2.7 Hz per volume. Fast z-plane shifts were controlled with a piezo collar. Visual stimuli were back-projected onto a vertically oriented 10 cm x 10 cm diffusive mylar screen positioned 5 cm from the fish using a pico projector (MP-CL1A, Sony). Stimuli subtended approximately 90° × 90° of visual angle, covering a large portion of the monocular visual field. Stimuli were rendered in OpenCV and presented using Bonsai^48,49^. A TTL signal sent from the microscope at the start of scanning was used to trigger an Arduino to synchronize imaging with visual stimulation. A Genie Nano M640 NIR camera (Teledyne DALSA), zoom lens (Navitar) and an IR long-pass filter (ThorLabs) were used to acquire tail tracking video at 450 Hz with a resolution of 640 × 480. Following functional imaging, fish were anesthetized in 1.5 mM tricaine and a high-resolution anatomical stack was acquired (2 µm z-spacing) encompassing a larger volume than the functional imaging planes.

Larvae were next processed for fluorescent in situ hybridization chain reaction (HCR), as described above, to establish *rln3a* and *penka* cell-type identities. The high-resolution two-photon volume acquired after functional imaging was next registered to the HCR confocal volume using Greedy (https://github.com/pyushkevich/greedy). Registration accuracy was confirmed by visual inspection, requiring unambiguous, one-to-one correspondence of individual nuclei across the two volumes at single-cell resolution; only cells meeting this criterion were included in downstream analysis. Functional imaging planes were motion corrected using CaImAn’s motion correction module prior to visual inspection^50^. FIJI was then used to extract fluorescence intensity values from nuclei in functional imaging planes that overlapped spatially with *rln3a* or *penka* HCR staining. dF/F traces from identified cells were clustered using k-means (SciPy, Python). The number of clusters (k) was selected using the elbow method, identifying the point of diminishing returns in within-cluster sum of squares as a function of k.

To determine the transmitter identity of *rln3a* and *penka*-expressing cells, separate larvae were processed identically using probe combinations of either *gad1b*/g*ad2* or *vglut2a* alongside *penka* or *rln3a*, and imaged on the same confocal system. All experiments were approved by the University of Toronto Local Animal Care Committee (protocol 20011546) and adhered to the governance of the Canadian Council on Animal Care.

*University of California, San Diego experiments (Figure 3):*

### Husbandry

All procedures were approved by the University of California, San Diego’s Institutional Animal Care and Use Committee (IACUC). We used larval zebrafish for all experiments in this study, tested between 6 and 8 days post fertilization (dpf). Animals were group housed in a 14-10-h light-dark cycle, temperature-controlled room and raised according to Zebrafish International Resource Center guidelines. All experiments were conducted during the light period (9:00-18:00). Larvae were fed rotifers twice daily from 5-14 dpf, and artemia thereafter.

### Two-photon imaging and behavior

Two-photon calcium imaging experiments were performed using 6-8 dpf

*Tg(elavl3:Hsa.H2B-GCaMP6s)^jf5^* larvae, as described previously^51^. Zebrafish were mounted dorsal side up in a thin layer of 3% low-melting point agarose (Invitrogen) in the lid of a 35-mm Petri dish (Fisher) and positioned under a stereomicroscope (Leica, S9D). Once solidified, agarose was removed from around the tail posterior to the swim bladder and around the nose and mouth with a fine scalpel. Fish were then habituated in the Petri dish for 20 minutes in E3 solution (5 mM NaCl, 169 μM KCl, 330 μM CaCl2, 161 μM MgSO4·7H2O) at room temperature before behavioral testing. All behavioral experiments were controlled with Bonsai packages. Behavioral videos were acquired from a camera mounted below the fish, at 121 Hz (FLIR Grasshopper, #33-534) with a 12X Zoom lens (ThorLabs, MVL12X12Z) and long-pass filter (ThorLabs, FBH850-10). The fish was illuminated with infrared (IR) light (ThorLabs, M850L3). Thermal stimuli were delivered using a 980-nm DPSS laser (Changchun New Industries Optoelectronics Technology, MDL-III-980) through a 400-μm, 0.48 NA optical fiber (Doric Lenses) positioned 1 mm from the fish’s nose. Stimuli were presented at three intensities (7, 28, and 65 mW), each delivered for 5 s, with stimulus order randomized across trials. Inter-stimulus intervals were randomized among 85, 90, or 95 s.

Imaging was performed on a ThorLabs Bergamo II multiphoton microscope using a Ti:Sapphire laser (MaiTai DeepSee, Spectra-Physics) tuned to 920 nm for excitation. GCaMP6s fluorescence was collected using a Nikon 16x 0.8 NA water-immersion objective. Neuronal activity was collected from a volume containing the PAG region using 8 planes spaced 7 µm apart at 512x512 pixel resolution, at a rate of ∼3 Hz per volume. Immediately after behavior and functional brain imaging, a structural stack was obtained at 860nm, with 1µm spacing and 25x frame averaging, starting 20µm above the first z-plane, ending 20µm below the last z-plane. Imaged larvae were euthanized in ice water, and fixed in 4% PFA. These larvae were processed for HCR to establish *rln3a* and *penka* cell-type identities using the same protocol described above. HCR-stained larvae were imaged on the same two-photon microscope used for functional imaging. Functional imaging planes were motion corrected using Suite2p prior to registration, and after cell extraction in Suite2p, neuron locations in each plane were assigned to their position in the anatomical volume. The fixed-brain stack was registered to the live-brain volume, and those transformations were applied to the HCR channels, using ANTs. The registration parameters were: [‘-d’, ‘3’,‘’--float’, ‘1’,’--interpolation’, ‘WelchWindowedSinc’, ‘--use-histogram-matching’, ‘0’, ‘-r’, ‘[fixed, moving, 1]’,’-t’, ‘Translation’ ‘-t’, ‘Rigid[0.1]’, ‘-m’, ‘MI[fixed, moving, 1, 32, regular, 0.25]’, ‘-c’, ‘[200x200x200x0,1e-8,10]’, ‘-f’, ‘12x8x4x2’, ‘-s’, ‘4x3x2x1’, ‘-t’, ‘Affine[0.1]’, ‘-m’, ‘MI[fixed, moving, 1, 32, regular, 0.25]’, ‘-c’, ‘[200x200x200x0,1e-8,10]’, ‘-f’, ‘12x8x4x2’, ‘-s’, ‘4x3x2x1’, ‘-t’, ‘SyN[0.05,3,0.1]’, ‘-m’, ‘CC[fixed, moving, 1, 2]’, ‘-c’, ‘[200x200x200x200x10,1e-7,10]’, ‘-f’, ‘12x8x4x2x1’, ‘-s’, ‘4x3x2x1x0’]. The high-resolution anatomical stack and post-HCR two-photon volume were checked with the same visual inspection criteria described above.

Behavioral time series of tail movements (>15 degrees) were cropped to the onset and offset of the 2-photon data acquisition periods. We excluded the first 5% of behavioral and neural measurements in each imaging experiment to avoid the influence of sound-evoked activity upon the initiation of scanning. Stimulus timing and calcium traces were aligned and resampled to a common 5 Hz sampling rate for further analysis. Each neuron’s fluorescence trace was detrended, smoothed with a 1-sec rolling mean, and dF/F traces were clustered using k-means as described above.

### Generation of relaxin-3a mutant line

Relaxin-3a knock out zebrafish were created using the CRISPR/Cas9 system^52^. Zebrafish relaxin-3a sequence was obtained from Ensembl (ENSDARG00000070780) and CRISPR targets were found in this sequence using the web-based tool ZiFit^53^. CRISPR targets that were upstream of the receptor-binding region were tested to produce a mutation to disrupt this region. The sgRNAs for these targets were produced using a method by Bassett et. al.^54^. To create the template, PCR was performed using Phusion High-Fidelity polymerase (ThermoScientific) with the 18-20bp target sequence (bolded) within the forward primer, 5’-GAA ATT AAT ACG ACT CAC TAT A**GG AGT AAA GGC GCT GGA CGC** GTT TTA GAG CTA GAA ATA GC-3’ and the universal reverse primer 5’-AAA AGC ACC GAC TCG GTG CCA CTT TTT CAA GTT GAT AAC GGA CTA GCC TTA TTT TAA CTT GCT ATT TCT AGC TCT AAA C-3’, which produced an approximately 100 bp product. The PCR reaction was cycled at (98°C 30s, 29 cycles of [98°C 10s, 60°C 30s, 72°C 15s], 72°C 10min, 10°C forever). The PCR product was purified with the QIAquick PCR purification kit (Qiagen). At least 200ng purified PCR template was used for transcription of sgRNA using the T7 MEGAscript transcription kit (Life Technologies) for 2-4 hours. After 1 hour of DNAse treatment, RNA was recovered using ammonium acetate precipitation at 4°C overnight and then eluted in 20 ul nuclease free water. Neat sgRNA solution (5 µg-10 µg) was aliquoted into 1µl portions and stored at -80°C for future use. To make the Cas9 mRNA, the expression vector pT3Ts-nCas9n (Addgene plasmid #46757) was linearized using XbaI and transcribed using the mMessage mMachine transcription kit (Life Technologies). Cas9 mRNA was then capped and polyadenylated using the Poly(A) kit (Ambion). After being precipitated in lithium chloride at 4°C overnight, Cas9 mRNA was eluted in 30 µl nuclease free water and stored similarly to the sgRNA. 1 µg of linearized plasmid template typically yielded about 900 ng-1.3 µg of mRNA. Zebrafish embryos were injected with 1nl of mixture containing 1 µl neat sgRNA with 1 µl neat Cas9 mRNA at the 1-cell stage.

### Detection of relaxin-3a mutants

To check for mutations in the *rln3a* locus, DNA was extracted from either 2-5 dpf embryos or adult tail fins by digesting in 20µg/ml Proteinase K solution in TE buffer at 55°C for 2 hours followed by denaturation of Proteinase K at 95°C for 10 mins. Primers were designed using VectorNTI which flanked the CRISPR target site, using the forward primer 5’-TCT CAG ACT TTA TCT CGC AGG GTT ATC A-3’ and reverse primer 5’-TCA AGC AAC TTA TAG AGC TTA GAG CTG AAA-3’, with a product size of 481bp. PCR was carried out using Taq polymerase (ThermoFisher Scientific) and cycled at (95°C 2mins, 29 cycles of [95°C 30s, 56.7°C 30s, 72°C 30s], 72°C 10min, 10°C forever). The mutated sequence was determined by sequencing the PCR product with BigDye (Life Technologies) and the forward primer. Wild type and the *rln3a^sq4sj^*mutant allele could be distinguished by using restriction fragment length polymorphism, as the restriction enzyme SfoI (NEB) cuts the mutant allele but not the wild type allele, giving two smaller bands at 247bp and 227bp. 4 µl of PCR product was used in a 10µl digest using SfoI and CutSmart buffer (NEB) and incubated at 37°C for 1 hour. The digested and undigested fragments corresponding to mutant or wild type allele could be resolved on a 1.5% agarose gel.

### Quantitative real time polymerase chain reaction (qRT-PCR)

Adult fish were euthanized in iced water then decapitated. Brains were dissected out in ice-cold Ringer’s saline (116mM NaCl, 2.9mM KCl, 1.8mM CaCl_2_, 5mM HEPES pH7.2). RNA was extracted using the RNeasy Mini Kit (Qiagen), where freshly dissected whole brains were kept in lysis buffer on ice until all brains were dissected, and then homogenized before proceeding with the manufacturer’s protocol. This extraction typically yielded about 100ng/µl RNA with A260/280 values between 1.80 and 2.10. 70ng of RNA was then reverse transcribed into cDNA using SuperScript III First Strand Synthesis System for RT-PCR (ThermoFisher), which yielded about 1µg of cDNA. Each reverse transcription reaction was run with a reverse transcriptase negative (RT-) control, to check for genomic DNA contamination. Primers were designed using the software Primer3 (Untergasser et al. 2012). The primers used were zebrafish β-actin1 forward 5’-AGA GCT ATG AGC TGC CTG ACG-3’ and reverse 5’ CCG CAA GAT TCC ATA CCC A 3’ (106 bp) as an internal control; and the gene-specific primer for relaxin-3a, forward 5’-CGT GCT GTC ATC TTC ACC TG-3’ and reverse 5’-GGG AGA TTT GAG TCA GTG GC-3’ (167 bp). Both primers spanned the exon-exon boundary, and were first verified by having a single peak on a post-PCR melt curve, which produced single bands after electrophoresis on agarose gel, with no bands with the RT-controls. qRT-PCR was run using Maxima SYBR Green/ROX qPCR Master Mix (ThermoFisher) on 1µl cDNA using the Applied Biosystems 7500HT Fast Real-Time PCR system. The PCR reaction was cycled at (95°C 10mins, 40 cycles of [95°C 15s, 60°C 1min], 95°C 10s, 60°C 15s, 10°C forever). 5 cDNA samples of each genotype (wild type and homozygous mutant) were run in duplicates, along with both the RT- and no template controls. Plotting a standard curve showed that the PCR reaction was between 80-110% efficiency, and hence the comparative C_t_ method ^43^ was used to analyze the group differences in expression. Mean ± s.d. and coefficient of variation (CV) for the relative expression (2^-ΔCt^) for each group was calculated as individual points as they were not paired samples. Fold change in expression was calculated by using the formula ΔCt = Ct_ref_ − Ct_gene_,^55^ and a 2-tailed unpaired Student’s T test with unequal variances was calculated to determine if the difference was statistically significant using Microsoft Excel.

### Chromogenic in situ hybridization

A DIG-labelled probe was generated against the entire coding region of the relaxin-3a gene using primers adapted from Donizetti et al^22^: forward 5’-AAA GCA CAG GTA GAC CAT CAG G-3’, and the reverse primer with a T3 start site at the 5’ end (bold) 5’-**AAT TAA CCC TCA CTA AAG GG** TGC AGC CCC ATT TGC AGC AGG-3’. 4dpf larval zebrafish in a nacre background were treated with 1-phenyl-2-thiourea (PTU) before fixing in 4% paraformaldehyde and stored in methanol at -20°C. Transcripts were then labelled using the Thisses’ protocol^56^.

Labeled larvae were cleared and mounted in glycerol. They were imaged using differential contrast microscopy, with a 20x 0.5 NA air objective on a Zeiss Imager M2 (Oberkochen, Germany) using an AxioCam Hrc camera. A z-stack, from the dorsal level of the brain to the interpeduncular nucleus, was collected with a step size of 1.075 µm and deconvolved using AutoDeblur (Media Cybernetics, Rockville, MD).

### Monitoring of locomotor activity

To monitor locomotor activity during the light-dark cycles, both mutants and wildtype siblings were put in a standard 48-well plate with wildtypes in the odd rows while mutants in the even rows from the top of the plate. Each well is 12 mm in diameter and filled with 1 mL fish water from the fish tank, and the 48-well plate was transferred into a light-proof and sound-proof incubator equipped with the behavior imaging setup. Images were taken at a rate of 5 Hz by a Basler camera above the 48-well plate with a lens (F2.2 f-25mm) and an IR filter (Midwest Optical Systems, longpass at 830 nm). Four IR LED bars (TMS Lite, length at 140 mm x 2 and 80 mm x 2) provided the IR light source for tracking the fish position during the dark phase. The IR LED bars with the same length are facing each other to prevent direct illumination to the fish. In between the IR LED bars and the 48-well plate, there is a semi-transparent plexiglass acrylic sheet (200 x 200 x 3 mm) serving as an insulator for heat generated from the IR LED bars and it can also homogenize the white light from below. The water temperature never surpassed 28.5°C in our setup. White light that serves as the circadian cue was provided by a 12V-powered light pad (ARTOGRAPH) and controlled by an Arduino board that was controlled by a custom-built real-time tracking program in Python.

### Visual-motor response

30-minute periods of darkness, alternating with 30-minute periods of light was provided. Statistical analysis for mutant behavior was performed using R Statistical Software (v4.4.1; R Core Team 2024). Total fish activity (distance moved in mm per minute) was measured over 60 minutes during the lighted habituation period or 30 minutes in each light/dark period in the two cycles. The data was divided into five epochs, based on a change in lighting, of baseline (habituation period), first dark/light cycle and second dark/light cycle. A linear mixed model was fitted to account for the repeated measures in the experimental design using the R package lmerTest v3.1.3^57^. Genotype, epoch and time in minutes were modelled as fixed effects on distance moved, while random effects of slope and intercepts for activity over time were estimated to model individual differences between subjects. An interaction term between genotype and epoch was included to test if genotype differences in distance moved depended on epoch. Type III F-tests with Kenward-Roger approximation were used to determine the fixed effects *p*-values. If the interaction effect was significant, pairwise contrasts between genotype groups within each epoch were extracted to determine the difference in activity between *rln3a* mutants and their wild-type siblings within each epoch. Satterthwaite’s approximation was used to estimate degrees of freedom for the post-hoc tests, with the Bonferroni correction applied to the *p*-values in this family of tests.

## Acknowledgements

This work was funded by the Ministry of Education of Singapore and the Biomedical Research Council of Singapore (SJ), Natural Sciences and Engineering Research Council of Canada, Canadian Foundation for Innovation, and Ontario Research Fund (TT) and the Klingenstein-Simons Fellowship in Neuroscience, the Sloan Research Fellowship, and NIH grant R00MH112840 (MLB). JC and SK were supported by scholarships from the NGS program at the National University of Singapore, and GMP was supported by an NSF GRFP.

---

**Supplemental Figure 1.**
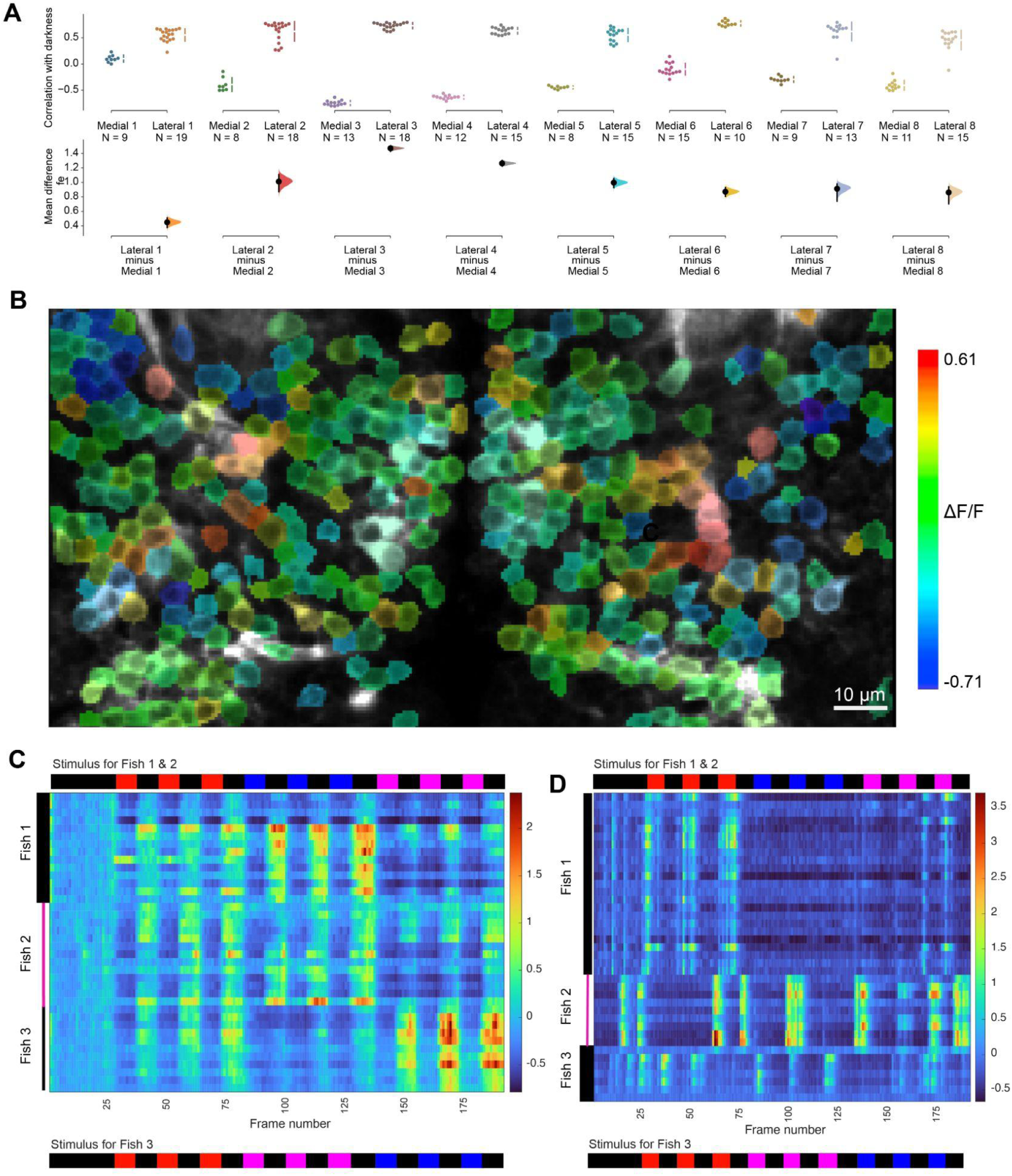
Cells in the lateral PAG show a consistent response to darkness. (**A**) Pearson coefficient of activity in PAG neurons, correlated with darkness, in 8 larvae exposed to 4 pulses of blue light, shown in a Cumming estimation plot^58^. The raw data is plotted on the upper axes; each mean difference is plotted on the lower axes as a bootstrap sampling distribution. Mean differences are depicted as dots; 95% confidence intervals are indicated by the ends of the vertical error bars. (**B**) Correlation map of GCaMP6f fluorescence change, relative to darkness. The fish was exposed to 3 pulses of red, blue and violet, as indicated at the top of panel C. (**C, D**). ΔF/F of neurons in the lateral (**C**) and medial (**D**) PAG of fish exposed to light pulses of different wavelengths. Fish 1 and 2 were exposed to the pattern shown above the plot, while fish 3 was exposed to the pattern shown below the plot.

**Supplemental Figure 2.**
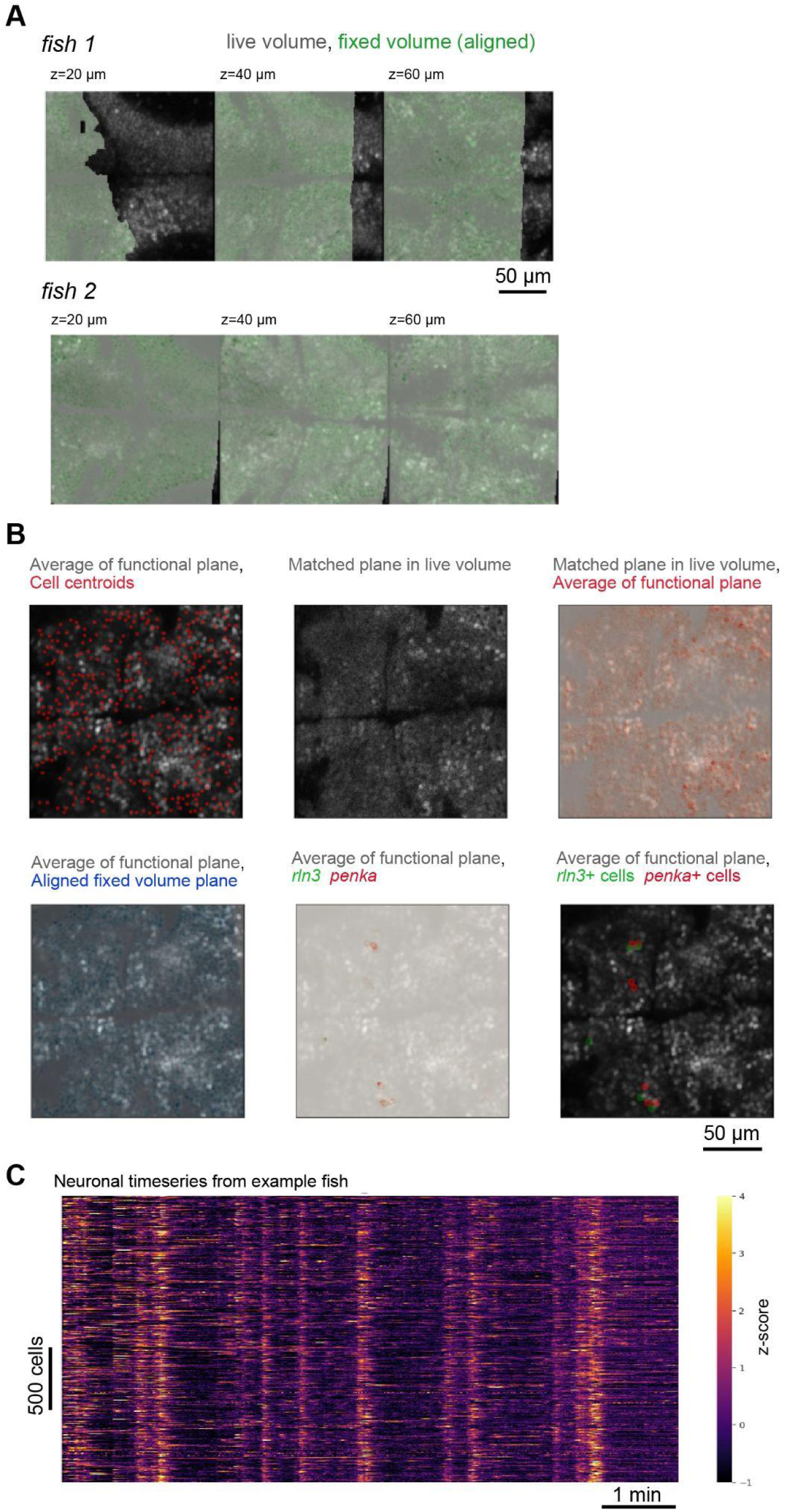
Registration of fixed brain volumes to live brain volumes, and assignment of cell identity. (**A**) Two example fish, showing alignment of post-registration fixed-brain GCaMP signal (green) on the live-brain GCaMP volume (grey), at 3 z-planes. (**B**) An example z-plane from an example fish, showing overlap of imaging data from functionally-recorded z-planes and cell identification, to live-brain volumes, to registered fixed-brain volumes (GCaMP and HCR signals). (**C**) Functional data in an example fish, of neural activity over time (z-scored fluorescence).

**Supplemental Figure 3.**
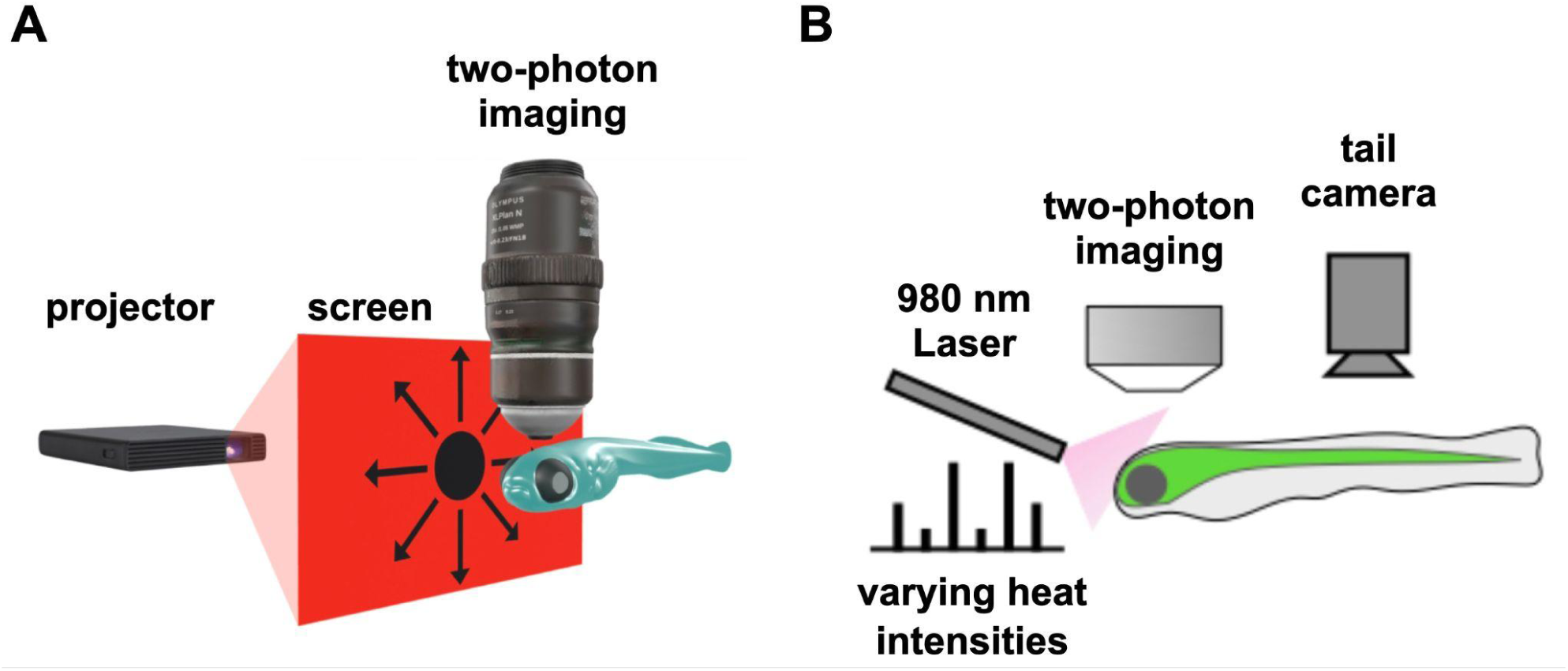
Experimental set up for stimulus delivery and imaging. (**A**) Loom stimulus. (**B**) Thermal stimulus.

**Supplemental Figure 4.**
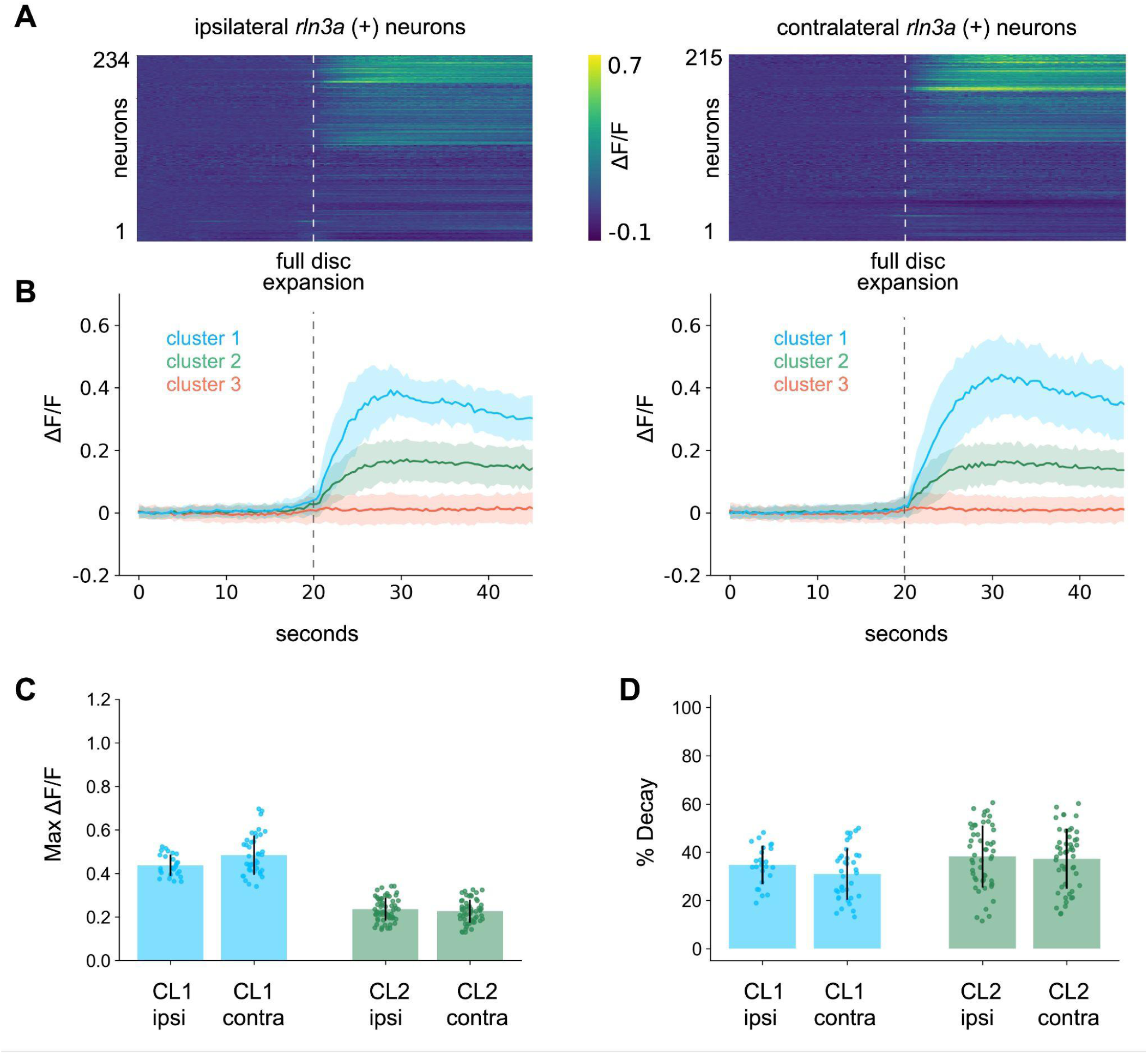
Darkness-evoked *rln3a* activity is bilaterally symmetric during monocular looming. (**A**) Heatmaps of ΔF/F for ipsilateral (left, n = 234) and contralateral (right, n = 215) *rln3a* neurons during monocular looming disc presentation, sorted by cluster assignment (n = 13 fish). Dashed line indicates full disc expansion at 20 s. (**B**) Mean ΔF/F ± SD per cluster for ipsilateral (left) and contralateral (right) populations. (**C**) Maximum ΔF/F for CL1 and CL2 neurons, ipsilateral and contralateral. (**D**) Percent decay for CL1 and CL2 neurons, ipsilateral and contralateral. Bars indicate mean ± SD. No significant differences were detected between ipsilateral and contralateral responses for maximum ΔF/F or percent decay (Wilcoxon rank-sum test).

**Supplemental Figure 5.**
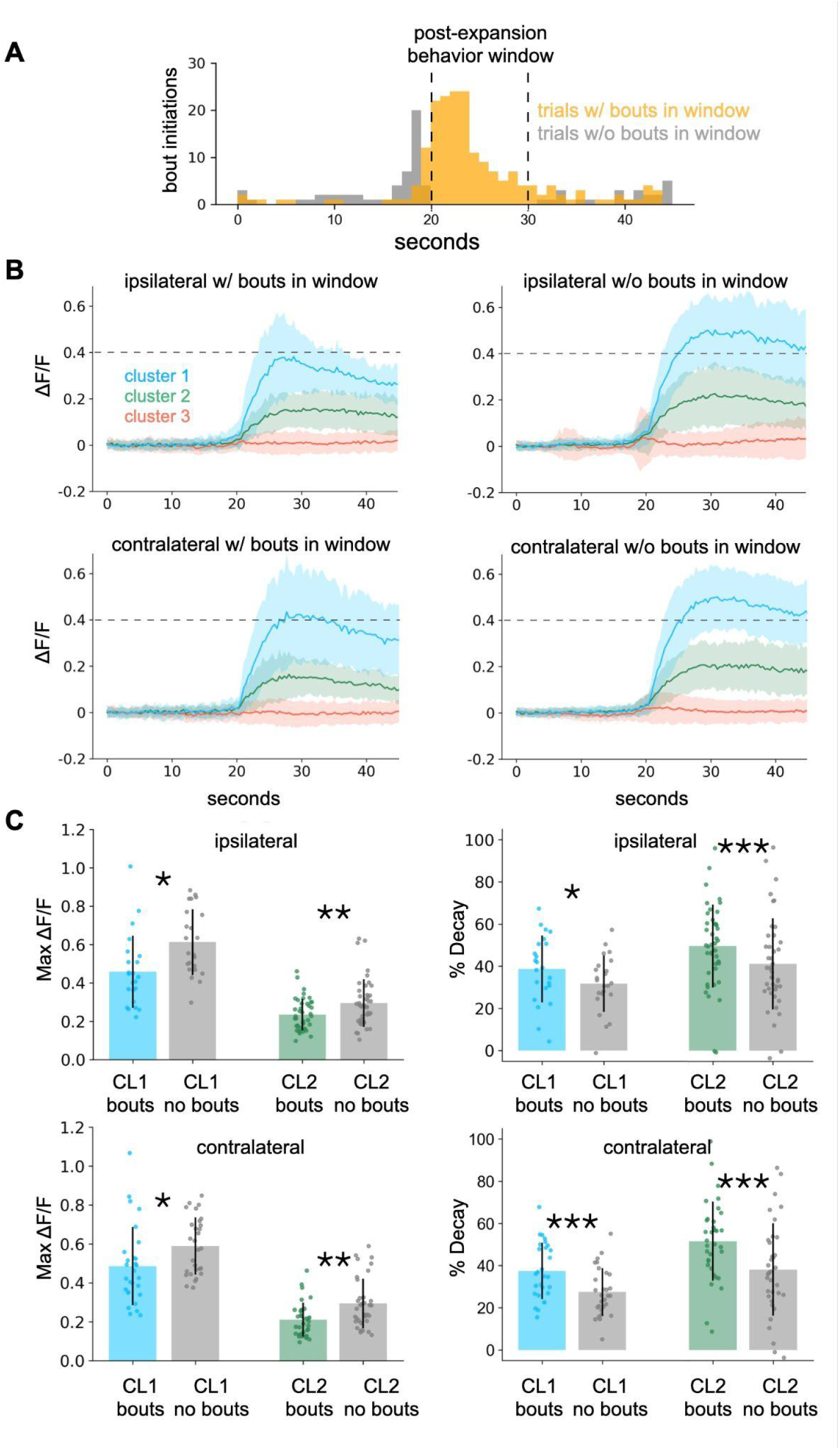
Higher *rln3a* activity is associated with post-loom behavioral suppression. **(A)** Bout initiations per 1-s bin during looming disc presentation, with trials split by presence (orange) or absence (gray) of swim bouts in the post-expansion behavioral window (dashed lines; n = 7 fish). **(B)** Mean ΔF/F ± SD for CL1 and CL2 *rln3a* neurons, ipsilateral and contralateral, for trials with versus without bouts in the behavioral window. **(C)** Maximum ΔF/F (left panels) and percent decay (right panels) for ipsilateral (top) and contralateral (bottom) CL1 and CL2 neurons on bout versus no-bout trials. Bars indicate mean ± SD. Asterisks indicate significance levels: \**p* < 0.05, \*\**p* < 0.01, \*\*\**p* < 0.001 (Wilcoxon signed-rank test).

**Supplemental Figure 6.**
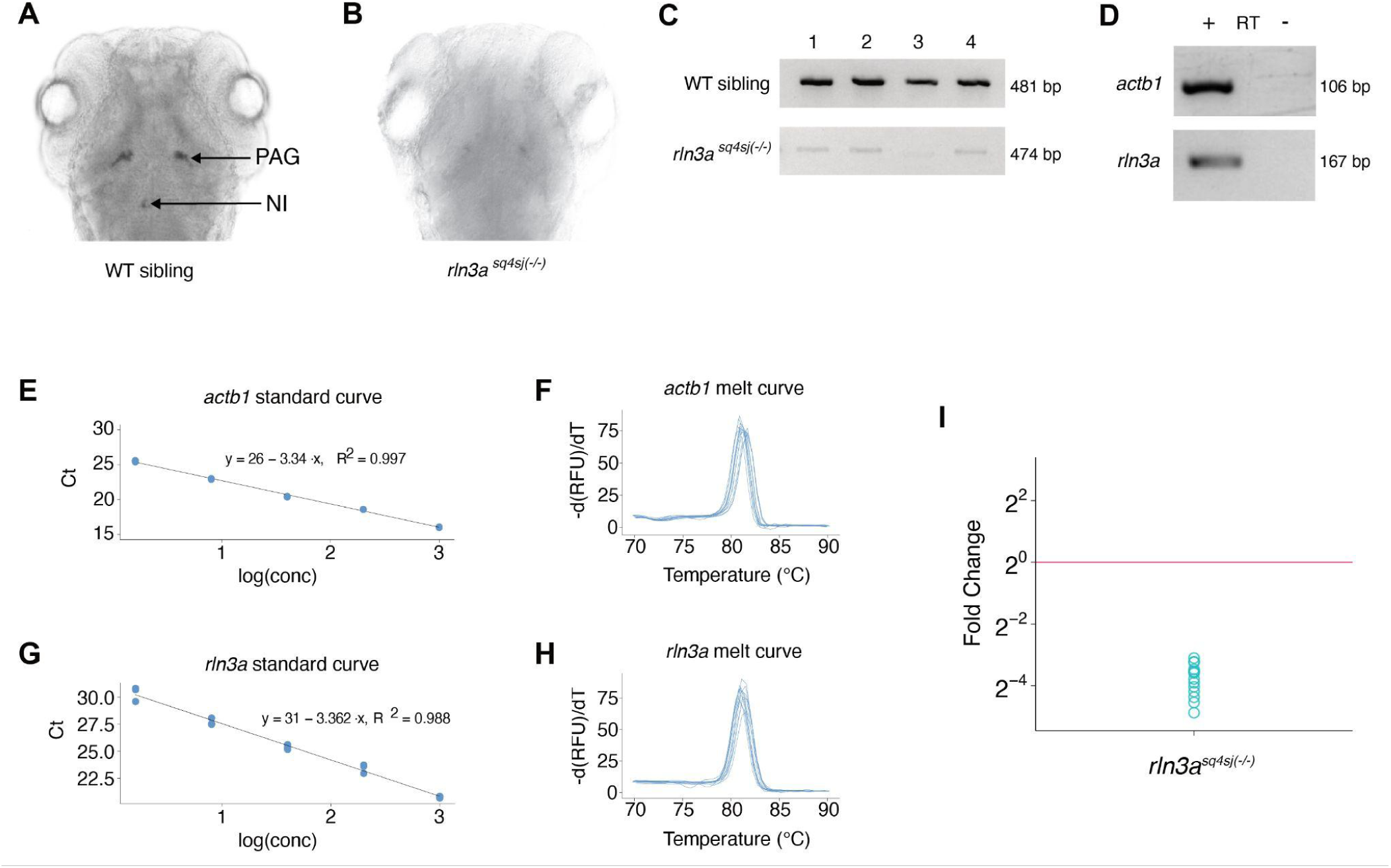
Molecular validation of the *rln3a^sq4sj^* mutant allele. **(A–B)** Wholemount ISH for *rln3a* in 4 dpf WT sibling (**A**) and *rln3a^sq4sj(-/-)^* (**B**) larvae showing strong PAG and NI signal in WT and near-absent signal in the mutant. **(C)** RT-PCR of full-length *rln3a* cDNA in four representative fish per genotype, showing a strong band in WT siblings (481 bp) and a faint, slightly smaller band in mutants (474 bp). **(D)** qRT-PCR products on agarose gel showing single bands for *actb1* (106 bp) and *rln3a* (167 bp), with no amplification in the RT− control. **(E–H)** Primer optimization: standard curves for *actb1* (E) and *rln3a* (G) showing ∼100% amplification efficiency; melt curves for *actb1* (**F**) and *rln3a* (H) showing single peaks confirming primer specificity. **(I)** Fold change of *rln3a* expression, with averaged WT expression set to 1 (pink line).

## Legends for Supplemental Movies

*Supplemental Movie 1*

Confocal z-stack of a 6 dpf larva showing *penka* (magenta) and *rln3a* (cyan) expression, stepping from dorsal to ventral. Anterior to the top.

*Supplemental Movie 2*

Transverse sections of the z-stack in Supplemental Movie 1, digitally resliced along the anterior-posterior axis, showing *penka* (magenta) and *rln3a* (cyan) expression. Sections step from anterior to posterior.

